# Turep: Detecting cross-cancer tumor-reactive T cells in single-cell and spatial transcriptomics data

**DOI:** 10.64898/2026.04.21.719961

**Authors:** Wendao Liu, Chia-Hao Tung, Eva M. Sevick-Muraca, Zhongming Zhao

**Affiliations:** The University of Texas MD Anderson Cancer Center UTHealth Houston Graduate School of Biomedical Sciences, Houston, TX, USA; Center for Precision Health, McWilliams School of Biomedical Informatics, The University of Texas Health Science Center at Houston, Houston, TX, USA; Center for Molecular Imaging, The Brown Foundation Institute of Molecular Medicine, The University of Texas Health Science Center at Houston, Houston, TX, USA

## Abstract

Tumor-infiltrating lymphocytes are essential for anti-tumor immunity, yet distinguishing tumor-reactive T cells from non-reactive bystander cells remains a significant challenge. Existing signatures, often derived from single cohorts, lack robustness in cross-cancer prediction. We present Turep, a deep learning method designed for robust, cross-cancer prediction of tumor-reactive T cells using single-cell or spatial transcriptomics data. By integrating paired single-cell RNA and T cell receptor sequencing data from seven human malignancies, we identified a pan-cancer tumor-reactive gene signature and leveraged generative data augmentation to address data imbalance. Turep consistently outperformed existing biomarkers, achieving a mean area under the receiver operating characteristic curve of 0.870 across cancer types. In validation across diverse cohorts, we found that Turep-predicted tumor-reactive T cell proportions could predict clinical response to immunotherapy. Furthermore, extending Turep to spatial transcriptomics revealed that tumor-reactive T cells preferentially resided in spatial niches where target cells exhibited elevated antigen presentation. Overall, Turep provides a powerful, generalizable tool for identifying tumor-reactive T cells and their spatial architectures, facilitating personalized cancer immunotherapy strategies.

## Introduction

The tumor microenvironment (TME) represents a highly complex and dynamic landscape where the interaction between malignant cells, stromal components, and immune infiltrates dictates the trajectory of cancer progression. Tumor-infiltrating lymphocytes (TILs), in particular CD8^+^ T cell populations are central to TME, acting as the primary effector for anti-tumor immunity^1,2^. Within the TME, these T cells are heterogeneous, consisting of a small number of tumor-reactive clones, which specifically recognize and respond to tumor-associated antigens or neoantigens, and a larger number of non-tumor-reactive bystander cells. Tumor-reactive T cells often transition into states of chronic activation and eventual exhaustion or differentiation into resident memory phenotypes, while bystander cells typically retain transcriptional profiles associated with viral specificities or homeostatic memory. Because the distinction between tumor-reactive and bystander T cells is lacking, this T cell heterogeneity often complicates the assessment of a patient’s functional immune status^3,4^.

The systematic identification of tumor-reactive populations has been significantly advanced by the advent of single-cell RNA sequencing (scRNA-seq) paired with single-cell T cell receptor (TCR) sequencing (scTCR-seq). This dual-omic approach allows for the high-resolution mapping of transcriptional phenotypes to specific TCR clonotypes, enabling researchers to link clonal expansion and functional states to antigen specificity. By integrating these technologies, it is possible to discern which T cell clones are actively engaged with known tumor antigens and which are merely occupying the niche without contributing to the anti-tumor response^5,6^.

To identify tumor-reactive T cells, antigen-centric strategies predict structural binding between major histocompatibility complex (MHC)-presented tumor epitopes and TCRs. These approaches require extensive prior knowledge of tumor epitopes. They are computationally extensive and frequently exhibit high false discovery rates when screened against large pools of potential epitopes. They may also fail to identify reactive TCRs if the candidate epitope pool is incomplete^7-9^.

Efforts to streamline the identification of tumor-reactive T cells using transcriptomics have led to the development of various gene signatures associated with tumor reactivity. So far, markers such as *CXCL13, GZMB*, and *ENTPD1*^10,11^, along with more comprehensive panels like NeoTCR8 and TR30^6,12^, have been proposed as indicators of tumor-reactive T cell states. However, existing signatures are optimized for specific cohorts and may face limitations in cross-cancer generalization due to the immense biological heterogeneity across different malignancies.

To overcome these challenges, we developed a deep-learning method, Tumor-reactivity prediction (Turep), for the cross-cancer transcriptome-based identification of tumor-infiltrating tumor-reactive T cells. It comprises of two components for prediction in scRNA-seq and spatial transcriptomics (ST) data respectively. By utilizing a generative data augmentation strategy and optimized loss functions, Turep addresses the inherent data imbalance between bystander and tumor-reactive T cells found in clinical samples, allowing for robust performance across multiple cancer indications.

The utility of Turep extends beyond standard single-cell analysis to the burgeoning field of ST. By applying Turep ST datasets, it becomes possible to not only estimate the proportion of tumor-reactive T cells in various therapeutic contexts but also to map their precise topographical distribution within the tissue. This capability allows for the investigation of spatial associations between reactive T cells and their target cells, providing critical insights into the physical interactions that drive clinical response to cancer immunotherapies.

## Results

### Tumor-reactive T cells phenotypes across cancer types

To systematically characterize and predict tumor-reactive T cells, we collected available paired scRNA-seq and scTCR-seq datasets across seven human malignancies (Supplementary Table S1): head and neck cancer (HNSC), breast cancer (BRCA), non-small cell lung cancer (NSCLC), gastrointestinal cancer (GIC), colon adenocarcinoma (COAD), rectal adenocarcinoma (READ), and skin cutaneous melanoma (SKCM)^5,6,13-15^. Tumor-reactive TCRs were experimentally validated using assays such as MANAFEST, ELISpot, and flow cytometry by being screened against identified tumor neoantigens^6,13^. Considering that CD8^+^ T cells are the primary effectors in tumor recognition, we focused our comparative analyses and predictive modeling on this lineage. We identified a pan-cancer tumor-reactive gene signature and developed a machine learning model by leveraging gene expression alone to predict reactivity across diverse indications. We further demonstrated the utility of this model by applying it to independent scRNA-seq and ST datasets to investigate the association between tumor-reactive T cells and therapeutic response, as well as to map active tumor-T cell niches (Fig. 1a).

**Fig. 1.**
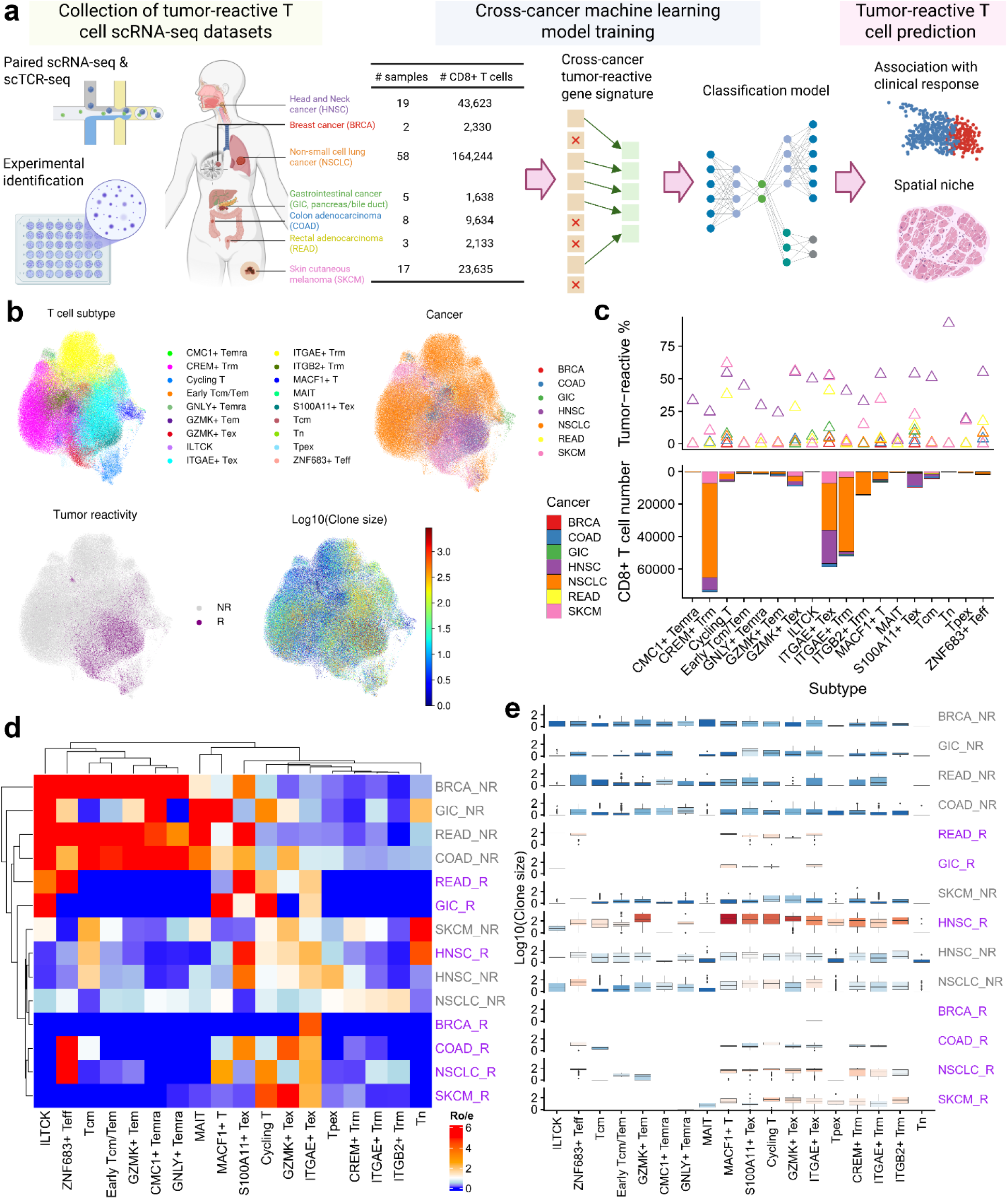
Characterization of tumor-reactive CD8^+^ T cell states across diverse malignancies. **a**. Schematic workflow illustrating the integration of paired scRNA-seq and scTCR-seq datasets, the identification of a pan-cancer tumor-reactive gene signature for machine learning model training, and subsequent model application to independent cohorts and spatial transcriptomics datasets. **b**. UMAP visualization of CD8^+^ T cells, colored by functional subpopulation, cancer type of origin, tumor reactivity status (R: reactive; NR: non-reactive), and log10-transformed clonal expansion size. **c**. Absolute cell counts (bottom) and the percentage of tumor-reactive T cells (top) within CD8^+^ T cell subpopulations across seven cancer types, highlighting the heterogeneity of reactive T cell abundance. **d**. Heatmap showing the enrichment of tumor-reactive and non-reactive T cells across functional states and cancer types, quantified by the ratio of observed to expected (Ro/e) values. **e**. Clonal expansion profiles (log10 clone size) of tumor-reactive and non-reactive T cells, stratified by subpopulation and cancer type, demonstrating the preferential expansion of reactive clones.

CD8^+^ T cell subpopulations were annotated using scRNA-seq profiles^16^, while tumor-reactivity status and clonal expansion levels were derived from scTCR-seq data (Fig. 1b). Across most cancer types, resident memory (Trm) and exhausted (Tex) T cells emerged as the most prevalent subpopulations, specifically CREM^+^, ITGAE^+^, and ITGB2^+^ Trm, alongside GZMK^+^, ITGAE^+^, and S100A11^+^ Tex (Fig. 1c). Although the proportion of tumor-reactive T cells within these subpopulations varied significantly across cohorts, likely reflecting a combination of biological heterogeneity and differences in experimental TCR identification methods, unique enrichment patterns were observed. Specifically, tumor-reactive cells were consistently enriched in exhausted (GZMK^+^, ITGAE^+^, S100A11^+^ Tex), ZNF683^+^ effector, and cycling subpopulations. This distribution likely reflected a natural T cell state transition from activation toward proliferation or exhaustion within an antigen-rich TME^17^. Conversely, bystander T cells were predominantly localized within memory states, including Trm, effector memory (Tem), and Tem re-expressing CD45RA (Temra, Fig. 1d).

We further evaluated the relationship between tumor reactivity and clonal expansion (Fig. 1e). Tumor-reactive T cells consistently exhibited high levels of clonal expansion across subpopulations and cancer types, whereas non-reactive T cells remained largely unexpanded. Given the established correlation between T cell expansion and patient outcomes^18-20^, these tumor-reactive clones likely represent a decisive factor in therapeutic efficacy. Notably, we also observed high clonal expansion in a subset of bystander T cells, suggesting the active recruitment and homing of these T cells into the tumor niche.

### A pan-cancer gene signature of tumor-reactive T cells

To identify conserved gene signatures of tumor-reactive T cells, we performed differential expression analysis between reactive and non-reactive populations within each cancer type (Supplementary Fig. S1, Table S2-S7). While only a small fraction of differentially expressed genes (DEGs) were universally shared across all cohorts, the majority exhibited either cancer-specific expression or were shared among a subset of malignancies (Fig. 2a, b).

**Fig. 2.**
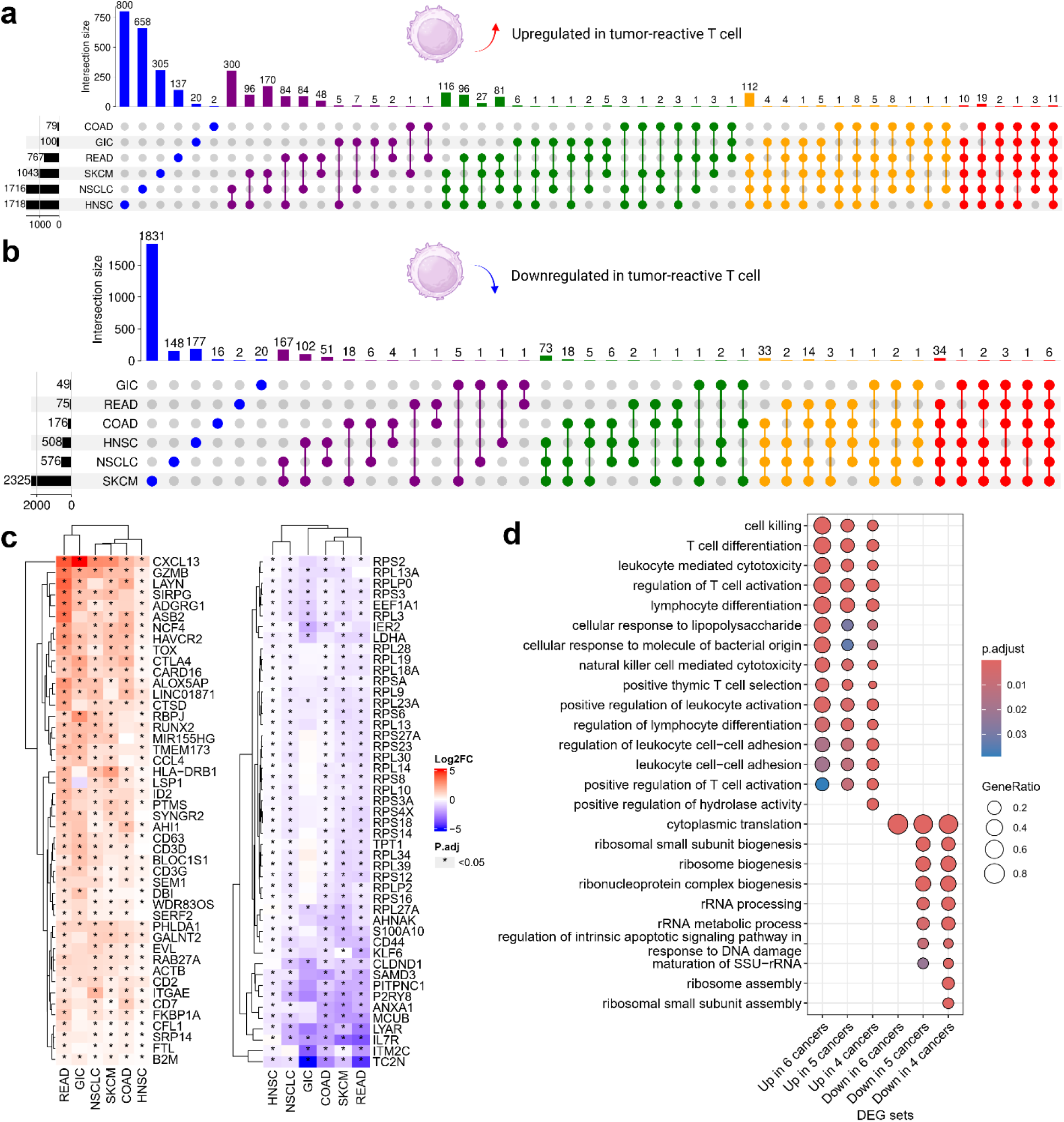
Identification of a pan-cancer transcriptional signature for tumor-reactive CD8^+^ T cells. **a, b**. UpSet plots illustrating the intersection of upregulated (**a**) and downregulated (**b**) differentially expressed genes (DEGs) across six human cancer types. Bar heights represent the number of genes unique to or shared between specific cancer cohorts. **c**. Heatmaps displaying the log2(Fold Change) of the most consistent pan-cancer DEGs (shared by at least five cancer types). Left: Upregulated genes associated with reactivity, including exhaustion and cytotoxicity markers (e.g., *CXCL13, GZMB, HAVCR2*). Right: Downregulated genes, highlighting metabolic and signaling shifts. **d**. Gene Ontology (GO) enrichment analysis of tumor-reactivity DEGs shared across four, five, or six cancer types.

We identified key drivers of T cell function and regulation among the genes upregulated in at least five cancers (Fig. 2c). Specifically, *CD3D, CD3G, CD2, ITGAE, CTLA4*, and *HAVCR2* emerged as central mediators of T cell activation and immune checkpoint regulation. Effector molecules including *GZMB, CCL4*, and *CXCL13* were also commonly upregulated. Furthermore, the upregulation of *ACTB* and *CFL1* linked to active cytoskeletal remodeling, possibly facilitating T cell migration and synapse formation within TME.

Conversely, the genes consistently downregulated across at least five cancers were dominated by ribosomal protein-encoding genes (e.g., *RPS2, RPL3*), a transcriptomic hallmark of T cell exhaustion. Other downregulated genes included *CD44*, involved in extracellular matrix (ECM) interactions; *IL7R*, a key component of homeostatic signaling; and *ANXA1*, which modulates T cell differentiation.

Pathway enrichment analysis further corroborated these findings (Fig. 2d). Conserved upregulated pathways were primarily associated with cell-mediated immunity, including T cell selection, activation, differentiation, adhesion, and cytotoxicity. Reflecting the observed downregulation of ribosomal genes, the corresponding pathways were significantly enriched for ribosome biogenesis, assembly, and rRNA processing, suggesting a metabolic shift or a specialized state of translational repression in tumor-reactive clones.

### Turep demonstrates superior performance in cross-cancer tumor-reactive T cell prediction

Leveraging the data from seven distinct malignancies, we developed Turep, a variational autoencoder (VAE)-based framework with additional classifiers for the robust, cross-cancer prediction of tumor-reactive T cells. To evaluate the model’s ability to generalize to unseen indications, we employed a leave-one-cancer-out cross-validation strategy, training the model on six cancer types and testing its performance on the remaining one (Fig. 3a).

**Fig. 3.**
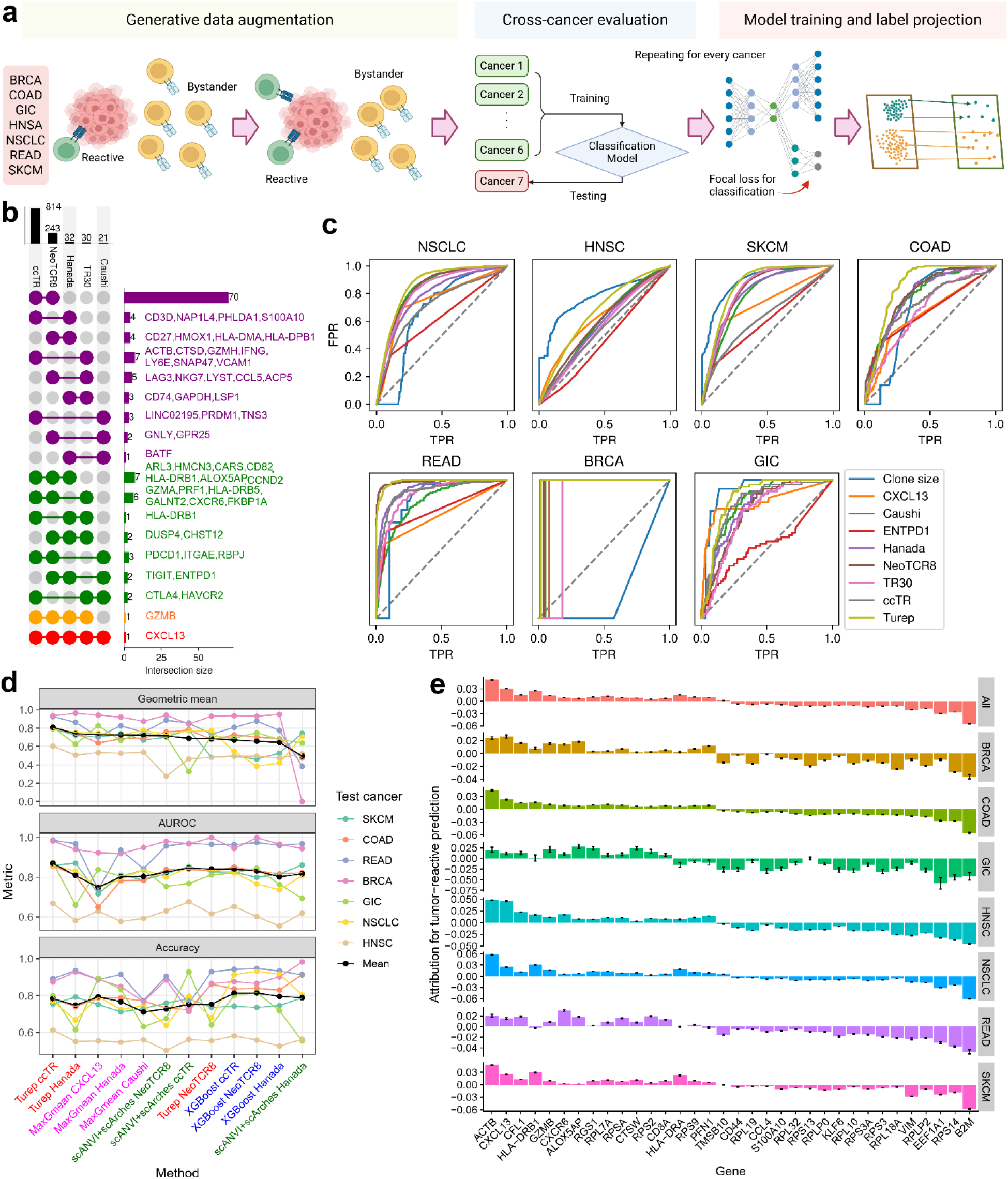
Turep for cross-cancer tumor-reactive T cell prediction. **a**. schematic diagram showing the process of generative data augmentation, cross-cancer leave-one-out training and evaluation, and final label projection onto scRNA-seq datasets. **b**. UpSet plot comparing the ccTR (cross-cancer tumor-reactive) gene signature with four previously reported T cell signatures (NeoTCR8, Hanada, TR30, and Caushi), highlighting both conserved and unique marker genes. **c**. Receiver Operating Characteristic (ROC) curves evaluating Turep’s predictive performance across seven cancer types. Turep is compared against single-gene biomarkers (e.g., *CXCL13*), established signatures, and clone size. **d**. Performance benchmarking of various machine learning architectures and signature-based methods across three metrics: G-mean, AUROC, and Accuracy. The top three models for each method are shown; full performance metrics are provided in Supplementary Table S9. **e**. Feature attribution analysis identifying the most influential genes for Turep’s classification. Attribution scores are shown for the global model (All) and individual cancer types, with error bars representing the standard error. Positive scores indicate features driving a “reactive” prediction.

To address the inherent class imbalance between tumor-reactive and bystander T cells within the datasets, we implemented a generative data augmentation pipeline. We upsampled the minority tumor-reactive population in three datasets and downsampled the majority non-reactive population in two large datasets (see Methods). Following this augmentation, the proportion of tumor-reactive T cells was stabilized at approximately 30% across all training sets, preventing the model from developing a classification bias toward the majority non-reactive class (Fig. 3a).

Architecturally, Turep builds upon the architecture from scANVI and scArches^21,22^, which were recently identified as top-performing methods for single-cell label transfer^23^. To further optimize performance on imbalanced biological data, we integrated focal loss into the classification objective, which compels the model to prioritize “hard-to-classify” cells during training^24^. The final Turep model utilizes an inclusive cross-cancer tumor-reactive (ccTR) signature of 814 genes, representing conserved patterns of up- and downregulation across the seven cancer types.

We benchmarked Turep against several established tumor-reactivity biomarkers (*CXCL13, ENTPD1*, and clone size) and gene signatures (NeoTCR8, TR30, Caushi, and Hanada, Supplementary Table S8)^6,10-13^.

Comparison of these signatures revealed that *CXCL13* was the only gene shared by all panels, reinforcing its central role in the tumor-reactive phenotype, while the cytotoxic marker *GZMB* was present in all but the Caushi signature (Fig. 3b).

We performed evaluation using receiver operating characteristic (ROC) curves. While individual markers or signatures perform well in specific contexts, they lack universal robustness (Fig. 3c). For instance, clone size proved highly predictive in HNSC, SKCM, and GIC, but exhibited poor performance in NSCLC and BRCA. Similarly, while the large NeoTCR8 panel was relatively robust, it failed to maintain high accuracy in HNSC. Remarkably, Turep consistently outperformed existing gene markers and signatures across the majority of cancer types, demonstrating its superior utility as a pan-cancer predictive tool.

Following the evaluation framework established by predicTCR^25^, which utilizes the geometric mean (G-mean) of sensitivity and specificity as a primary metric, we benchmarked Turep against several high-performing baseline models for cross-cancer prediction. These baselines included: XGBoost: The machine learning architecture employed by predicTCR; scANVI+scArches: Currently regarded as the gold standard for single-cell label projection; MaxGmean: A grid-search strategy designed to identify the optimal classification threshold that maximizes the G-mean for specific gene markers or signatures (Fig. 3d). For each architecture, we systematically tested various gene signatures as input features to identify the optimal model configuration (Supplementary Table S9). Turep, integrated with our identified ccTR signature, achieved the best performance across all cohorts, with a mean G-mean of 0.810 and a mean area under receiver operating characteristic curve (AUROC) of 0.870. While other established signatures as input features also performed well within these machine learning frameworks, they did not reach this performance. For example, Turep utilizing the Hanada signature achieved a G-mean of 0.740 (AUROC 0.808), marginally outperforming the MaxGmean model using the same signature (G-mean 0.727; AUROC 0.803). Models trained on original data without generative data augmentation generally had lower G-mean and AUROC values (Supplementary Fig. S2).

To investigate the molecular logic underlying Turep’s predictions, we calculated SHapley Additive exPlanations (SHAP) values to identify the most influential features for classification (Fig. 3e)^26^. Several genes emerged as highly influential across all cancer types, including *CXCL13, ACTB, CFL1, GZMB*, and *CD8A*, suggesting a conserved core program of tumor reactivity. However, we also observed significant inter-cancer variability in feature importance for genes such as *HLA-DRA* and *HLA-DRB1*.

Interestingly, we noted that many prominent DEGs identified in our initial analysis exhibited differing levels of importance in the classification model (Fig. 2). This divergence likely stems from our use of generative data augmentation and focal loss, which compel the model to learn complex, non-linear patterns that differentiate “hard-to-classify” tumor-reactive cells from bystanders, rather than relying solely on the most abundant transcriptional differences.

### Turep-predicted tumor-reactive T cells are strongly associated with clinical response

Given the pivotal role of T cells in anti-tumor immunity, transcriptomic signatures of reactivity are increasingly recognized as correlates of immunotherapy efficacy. Unlike aggregated gene expression scores, the Turep model provides a granular estimation of the actual proportion of tumor-reactive CD8^+^ T cells in tumor-infiltrating CD8^+^ T within any given sample. This sample-level metric offers a more precise readout of TME. For instance, a predicted proportion near zero suggests a tumor niche lacking endogenous reactivity, potentially indicating a need for interventions such as cancer vaccines or CAR-T cell therapy to de novo prime or engineer an immune response.

We evaluated the association between predicted reactivity and clinical outcomes across five diverse treatment datasets. In the BioKey BRCA cohort, which includes longitudinal samples from patients undergoing anti-PD1 and chemotherapy, we observed a striking difference between patients based on T cell clonal expansion—a known proxy for therapeutic response^27^. Predicted tumor-reactive T cells were virtually absent in the non-expanded (NE) group, whereas the expanded (E) group exhibited significantly higher proportions (Fig. 4a). Interestingly, anti-PD1 treatment did not markedly shift these proportions in most patients. Furthermore, patients receiving concurrent chemotherapy showed a slight reduction in tumor-reactive T cell abundance compared to those receiving immunotherapy alone, likely reflecting the suppressive effect of chemotherapy on T cell proliferation (Fig. 4a). These results were corroborated in the GSE246613 BRCA dataset^28^, where responders (R1 and R2 groups) consistently harbored a higher frequency of predicted tumor-reactive clones compared to non-responders (NR) (Fig. 4b).

**Fig. 4.**
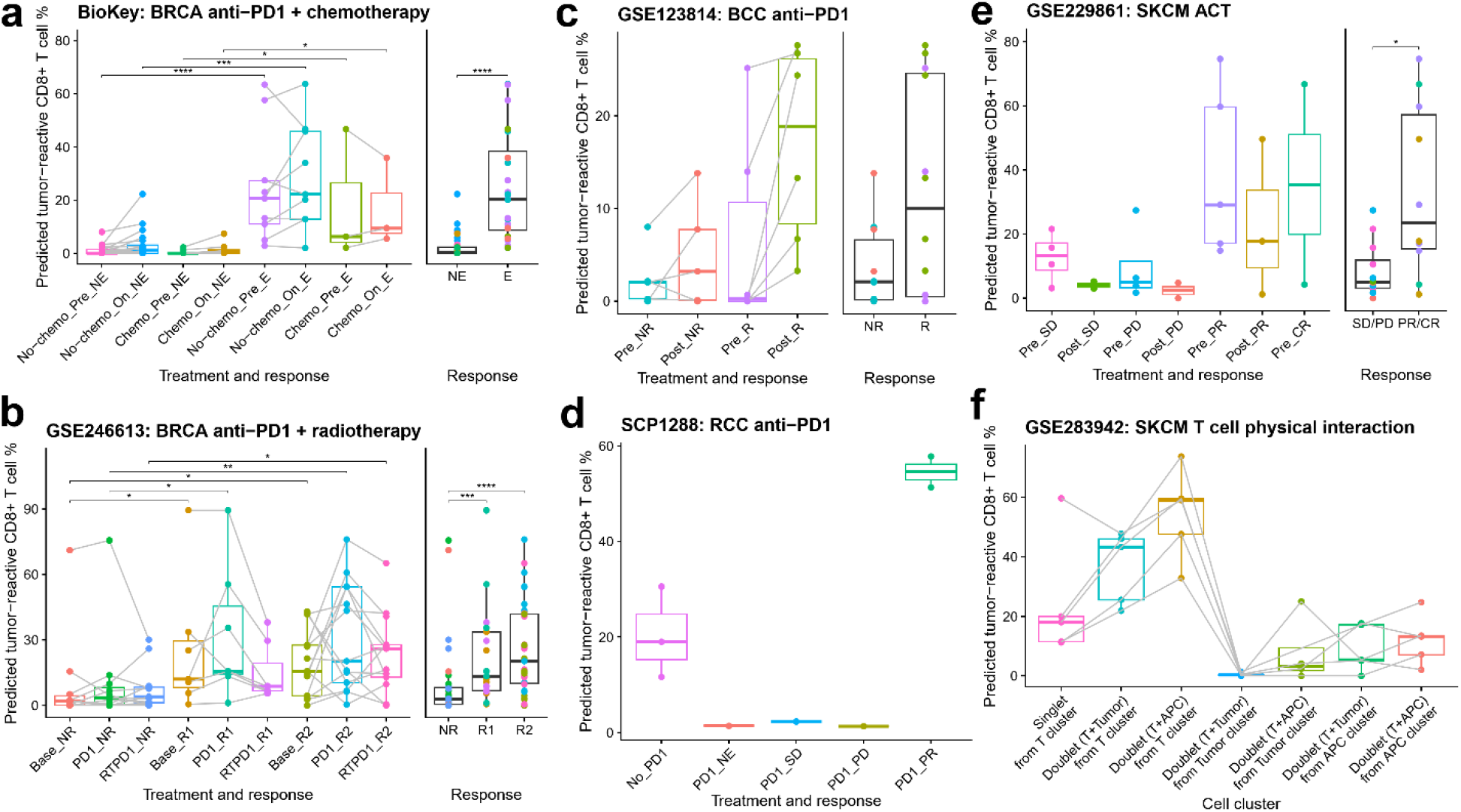
Turep tumor-reactive CD8^+^ T cell prediction acts as a biomarker for clinical response. **a**. Predicted percentage of tumor-reactive CD8^+^ T cells in the BioKey breast cancer (BRCA) cohort, stratified by chemotherapy treatment, anti-PD1 therapy status (Pre- or On-therapy), and T cell clonal expansion (E: expanded; NE: non-expanded). **b**. Predicted tumor-reactive CD8^+^ T cell abundance in the GSE246613 BRCA cohort across treatment stages: Baseline (Base), post-anti-PD1 (PD1), and post-combination radiotherapy and anti-PD1 (RTPD1). Patients are grouped by clinical response (R1/2: responders; NR: non-responders). **c**. Predicted reactivity percentages in basal cell carcinoma (BCC; GSE123814) before (Pre) and after (Post) anti-PD1 therapy, categorized by patient response (R: responder; NR: non-responder). **d**. Abundance of tumor-reactive T cells in renal cell carcinoma (RCC; SCP1288) samples across clinical response categories: not evaluable (NE), stable disease (SD), progressive disease (PD), and partial response (PR). **e**. Predicted reactivity in melanoma (SKCM; GSE229861) following adoptive cell transfer (ACT). Samples are categorized by timepoint (Pre-vs. Post-ACT) and clinical outcome (SD/PD vs. PR/CR). CR: complete response. **f**. Analysis of predicted tumor-reactive T cells in melanoma (SKCM; GSE283942) focusing on multi-cellular doublets. Data illustrate reactivity in singlets vs. doublets captured from T cell, tumor, or antigen-presenting cell (APC) clusters, indicating physical interaction niches. Statistical significance was determined by Wilcoxon rank-sum test: *: P < 0.05, **: P < 0.01, ***: P < 0.001, ****: P < 0.0001.

To test the model’s robustness, we applied Turep to two malignancies not included in the original training data. In the first basal cell carcinoma (BCC) cohort (GSE123814)^29^, responders not only exhibited higher baseline reactivity but also showed a significant elevation in predicted tumor-reactive T cell proportions following anti-PD1 therapy (Fig. 4c). This suggests a therapy-induced infiltrating or localized proliferation of reactive clones. Similarly, in the second renal cell carcinoma (RCC) dataset (SCP1288)^30^, patients with stable (SD) or progressive disease (PD) lacked detectable tumor-reactive T cells, while those with a partial response (PR) displayed high predicted tumor-reactive proportions (Fig. 4d).

The utility of Turep extends beyond immune checkpoint blockade to adoptive cell transfer (ACT). In the GSE229861 TIL-ACT dataset^31^, clinical benefit (PR/CR) was strongly associated with a high proportion of tumor-reactive T cells (Fig. 4e). Notably, the ACT process itself—which involves massive *ex vivo* expansion of both tumor-reactive and bystander T cells without selection—did not increase the proportion of tumor-reactive cells relative to the total T cell pool, suggesting that its primary therapeutic effect is driven by increasing absolute cell numbers rather than shifting the clonal composition. This implies that non-responders may require specialized TCR clonotype selection or TCR/CAR-engineering to enrich the infusion product with reactive clonotypes.

Finally, we demonstrated the physical interactions between predicted tumor-reactive T cells and target cells using a recent study about heterotypic clusters of CD8^+^ T cells and tumor cells or antigen-presenting cells (APCs) ^32^. In scRNA-seq data, these physical interactions are often captured as doublets or multiplets. Using Turep, we confirmed that while T cell singlets generally lacked tumor-reactive clones, doublets containing CD8^+^ T cells and tumor cells or APCs were significantly enriched for predicted reactivity (Fig. 4f). This finding suggests that Turep-predicted cells are not only transcriptionally reactive but also more likely to be in functional physical contact with their targets in the tumor niche.

### Spatial proximity and interaction niches of tumor-reactive T cells

High-resolution ST platforms enable single-cell resolution profiling, allowing for the precise identification of tumor-reactive T cells and the exploration of their spatial architecture. To locate these cells, we utilized the SPATCH dataset, which systematically profiled three cancer types: COAD, hepatocellular carcinoma (HCC), and ovarian cancer (OV) across four distinct ST technologies^33^. Considering the inherent challenges in cross-technology integration, we developed Turep-st, an adaptation of the FADVI framework^34^. This model disentangles label-associated biological variation from technology-specific batch effects into distinct latent spaces, facilitating the seamless projection of reactivity labels from scRNA-seq references onto ST query data (Fig. 5a, Supplementary Fig. S3).

**Fig. 5.**
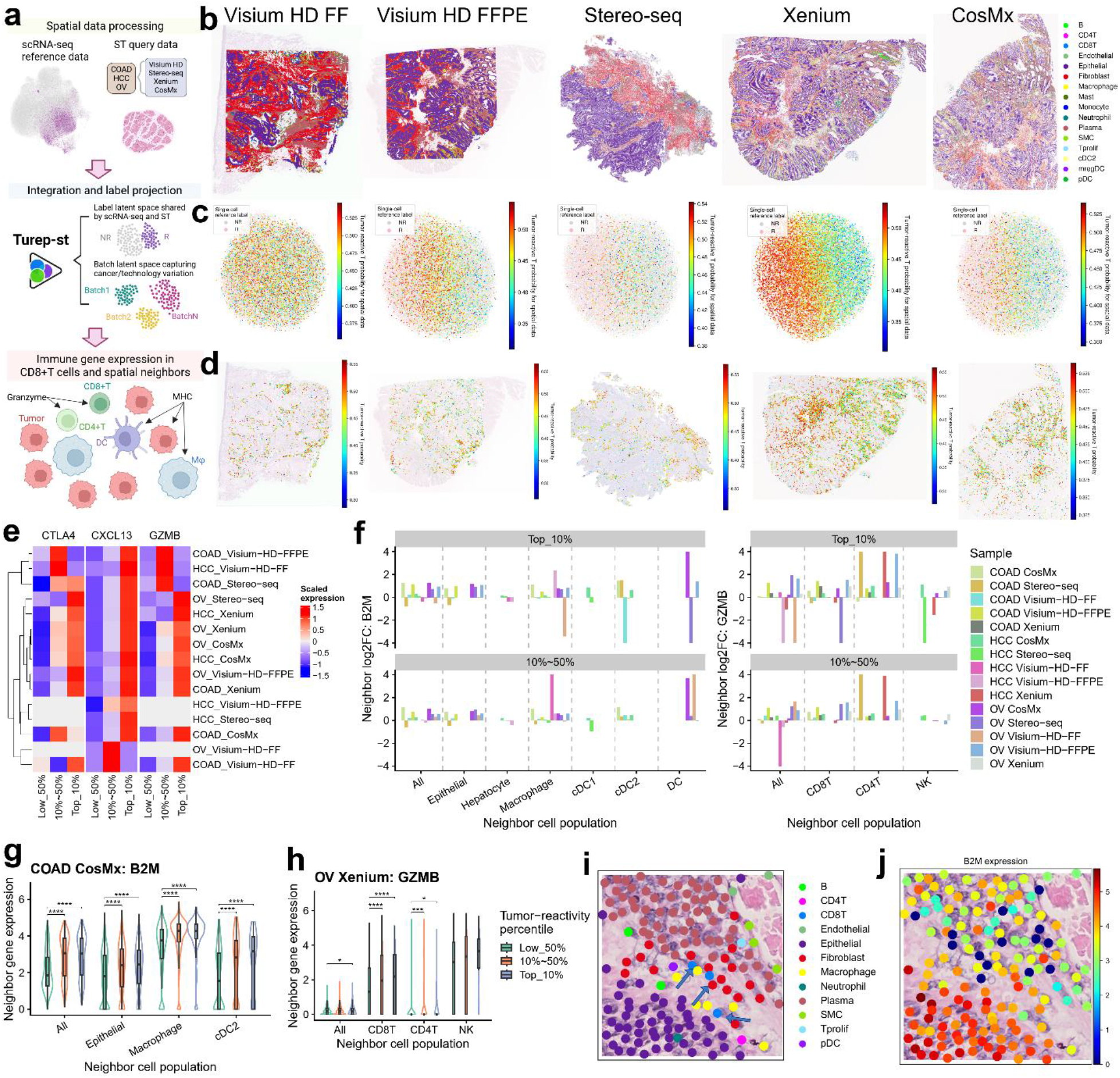
Spatially resolved identification and niche analysis of tumor-reactive T cells. **a**. Schematic workflow of the Turep-st framework, illustrating the integration of scRNA-seq reference data with diverse ST platforms (Visium HD, Stereo-seq, Xenium, CosMx) for label projection and subsequent spatial neighborhood analysis. FF: fresh frozen; FFPE: formalin-fixed paraffin-embedded. **b**. Spatial map of cell-type distributions in a colon adenocarcinoma (COAD) sample, with labels overlaid on the corresponding hematoxylin and eosin (H&E) stained tissue section. **c**. UMAP visualization of the shared latent space between scRNA-seq reference and ST query data, colored by predicted tumor-reactivity probability. Turep-st utilizes a joint embedding to facilitate robust cross-platform prediction. **d**. Spatial visualization of predicted tumor-reactivity probabilities for CD8^+^ T cells, mapped back onto the original tissue sample. **e**. Scaled expression of *CTLA4, CXCL13, GZMB* across T cells grouped by predicted reactivity. **f**. Differential expression analysis of *B2M* and *GZMB* in neighboring cell populations. Bar plots represent the log2(Fold Change) in neighbors of potentially tumor-reactive T cells (top 10% or 10%∼50% reactivity percentile) compared to non-tumor-reactive T cells (bottom 50% reactivity percentile) across multiple ST datasets and cancer types. **g**. Violin plots showing elevated *B2M* expression in epithelial and macrophage neighbors of potentially reactive T cells in the COAD CosMx dataset. **h**. Violin plots showing elevated *GZMB* expression in CD8^+^ and CD4^+^ T cell neighbors of potentially reactive T cells in the OV Xenium dataset. Statistical significance was determined by Wilcoxon rank-sum test: *: P < 0.05, **: P < 0.01, ***: P < 0.001, ****: P < 0.0001. **i-j**. Representative images of active tumor-recognition region in COAD. **i**, Spatial proximity between predicted tumor-reactive T cells (blue arrows) and specific cell types, showing reactive T cells localized near B2M^high^ tumor cells and APCs. **j**, Corresponding *B2M* expression levels in this region.

In COAD ST samples, where epithelial tumor cells and fibroblasts constitute the dominant populations (Fig. 5b), we isolated the CD8^+^ T cell compartment for joint embedding with our reference data. Turep-st utilized the shared latent subspace to calculate tumor-reactivity probabilities. T cells similar as reference tumor-reactive cells were assigned high probabilities, while those aligned with bystander cells were assigned low probabilities (Fig. 5c). Spatial mapping revealed that tumor-reactive and non-reactive T cells are often sequestered in distinct topographical regions within TME (Fig. 5d).

Building on our observation that tumor-reactive T cells are enriched in heterotypic clusters (Fig. 4f), we investigated whether predicted tumor-reactive T cells maintain closer physical proximity to target cells. We defined the top 10% and top 50% predicted reactivity percentiles as high-confidence and potentially tumor-reactive T cell populations, respectively. These T cells consistently showed high expression of genes associated with tumor reactivity, including *CXCL13, GZMB*, and *CTLA4* (Fig. 5e). While the overall neighbor composition varied across ST platforms due to technical differences in capture and resolution (Supplementary Fig. S4), a consistent functional pattern emerged regarding antigen presentation. Because MHC-I is essential for CD8^+^ T cell recognition, we quantified the expression of *B2M*—a necessary component for the MHC-I complex—in the spatial neighbors of T cells. We focused on the *B2M* gene because most HLA genes were not profiled by ST technologies, and *B2M* gene expression is regulated together with HLA genes^35^. Across all ST samples and platforms, neighbor *B2M* expression was generally higher for T cells in the top 10% and 10∼50% reactivity groups (Fig. 5f). Furthermore, neighbors involved in cell-mediated immunity exhibited significantly elevated granzyme levels, including *GZMB, GZMA*, and *GZMK* (Fig. 5f, Supplementary Fig. S5). For instance, in the COAD CosMx dataset, epithelial cells, macrophages, and cDC2s neighboring reactive T cells showed significantly upregulated *B2M* compared to neighbors of T cells in the bottom 50% percentile (Fig. 5g). Similarly, in the OV Xenium dataset, neighboring CD8^+^ and CD4^+^ T cells exhibited significantly higher *GZMB* expression in reactive niches (Fig. 5h).

By integrating predicted reactivity probabilities with neighbor transcriptomics, we can effectively map active tumor-recognition regions. As illustrated in the COAD CosMx sample (Fig. 5i, j), we identified localized niches where high-probability tumor-reactive T cells are found in close physical contact with B2M^high^ epithelial tumor cells and APC populations. This represents the spatial manifestation of the anti-tumor response, highlighting specific regions of active immune engagement and antigen sensing within the complex tissue architecture, which closely resemble the reported cancer regions of antigen presentation and T cell engagement and retention (CRATER)^36^. This spatial validation confirms that Turep-st not only identifies transcriptionally tumor-reactive cells but correctly locates them within functionally relevant, antigen-presenting neighborhoods.

## Discussion

In this study, we developed Turep, a deep learning-based framework for the systematic identification and spatial mapping of tumor-reactive T cells across diverse human malignancies. By integrating paired scRNA-seq and scTCR-seq data from seven cancer types, we defined a conserved ccTR signature and demonstrated that a generative data augmentation strategy combined with a focal loss objective significantly improves classification accuracy. Our results demonstrated that Turep could effectively identify tumor-reactive clones across both single-cell and spatial transcriptomics platforms, providing a robust tool for investigating the functional architecture of TME.

The primary strength of Turep lies in its superior generalization capability across different cancer types. Unlike previous marker-based or signature-based approaches that often exhibit variable performance^12^, Turep consistently maintained high predictive accuracy even when applied to unseen cancer types. This robustness is achieved through as an inclusive pan-cancer signature that captures conserved programs of exhaustion and activation, as well the disentanglement of biological signals from batch effects. By outperforming competing models like XGBoost and scANVI in leave-one-cancer-out cross-validation, Turep establishes a new benchmark for antigen-agnostic T cell reactivity prediction.

Despite its robust performance, several limitations of this study warrant consideration. First, the training data was limited to seven cancer types due to the limitation of available datasets; while we demonstrated successful generalization to BCC and RCC, a larger and more diverse training set encompassing more malignancies would likely further enhance the model’s reliability. Second, the validation of tumor reactivity in spatial transcriptomics remains challenging due to the current lack of gold-standard, ground-truth labels for TCR specificity in spatial data. While we used spatial proximity to cells and heterotypic doublets as high-confidence proxies for reactivity, direct experimental validation of TCR-antigen interactions in a spatial context would be necessary to definitively confirm these interaction niches.

The potential clinical applications for Turep are broad and impactful. As a predictive biomarker, the model can be used to stratify patients for immunotherapy response by quantifying the baseline proportion of tumor-reactive T cells, potentially identifying non-responsive tumors that require specialized priming strategies such as cancer vaccines. In the realm of ACT, Turep could serve as a screening tool to enrich TIL products for tumor-reactive clonotypes during *ex vivo* expansion, potentially improving therapeutic efficacy. Furthermore, by identifying potential tumor-reactive TCRs, our findings could inform the design of CAR-T therapies or synthetic receptors optimized for survival and function.

Collectively, our study demonstrates that the transcriptomic state of tumor-reactive T cells is conserve dacross cancers to permit accurate prediction using machine learning. Turep offers a scalable and versatile framework for characterizing the anti-tumor immune response in high resolution. As single-cell and spatial technologies become increasingly integrated into clinical diagnostics, tools like Turep will be essential for translating complex multi-omics datasets into actionable insights for personalized cancer immunotherapy.

## Methods

### Data collection and processing

We used five paired scRNA-seq and scTCR-seq datasets covering seven cancer types to identify cross-cancer tumor-reactive T cell signatures and train our machine learning model. These included GEO datasets GSE176021 and GSE180268^13,14^, as well as dbGaP datasets phs001451, phs002748, and phs002765^5,6,15^. All data came from tumor-infiltrating T cells. For GSE176021, GSE180268, and phs002765, we used the processed gene expression matrices and TCR clonotype tables provided by the authors. For phs001451 and phs002748, we processed the raw sequencing files using Cellranger-multi (v7.0.0). Detailed information for all datasets is available in Supplementary Table S1.

We identified tumor-reactive T cells based on their TCR clonotypes. T cells experimentally validated with tumor-specific TCRs against tumor neoantigens in the original studies were labeled as “tumor-reactive”. All other cells were labeled as “non-tumor-reactive/bystander”.

### scRNA-seq and scTCR-seq data processing

We used Scanpy (v1.11.4) and Scirpy (v0.19.0) to process and analyze paired scRNA-seq and scTCR-seq data^37,38^. Quality control was performance by removing cells with low number of expressed genes and high mitochondrial count percentage in scRNA-seq data. We did additional filtering by removing T cells without detected TCR information. The filtered data were then normalized to 10,000 total library size and log-transformed.

Most previous studies only identified CD8^+^ TCR clonotypes specific to MHC-I complexes. Because CD4^+^ tumor-specific TCRs were rarely identified, we separated the CD8^+^ and CD4^+^ T cells and focused on the CD8^+^ population. For each dataset, we selected 2,000 highly variable genes for principal component analysis (PCA) and unsupervised Leiden clustering. Clusters with high *CD8A* expression were kept for downstream analysis and model training.

For scTCR-seq data, we defined clonotypes using both the α and β chain amino acid sequences. If a cell contained dual TCRs, we only considered the most abundant pair of chains. We defined clone size as the number of T cells in a sample sharing the same TCR clonotype.

We used scAtlasVAE (v1.0.5.9) for annotation of CD8^+^ T cell subpopulations^16^. The model is pretrained on a large collection of CD8^+^ T cells from multiple diseases including cancers. Model “huARdb_v2_GEX.CD8.hvg4k.supervised.model” and reference dataset

“huARdb_v2_GEX.CD8.hvg4k.h5ad” were used to transfer the reference T cell subpopulation labels to the query datasets.

### Subpopulation enrichment of tumor-reactive T cells

To identify which T cell subpopulations were enriched for tumor reactivity, we calculated the ratio of observed to expected (Ro/e) cell numbers for each subset.

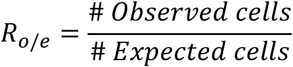

We used a Chi-square test to determine the expected cell counts for each subpopulation within each cancer type and reactivity group. Subpopulations with an *R*_*o*/*e*_>1 were considered enriched for either tumor-reactive or non-tumor-reactive T cells.

### Gene signature analysis

We identified DEGs between tumor-reactive and non-tumor-reactive T cells for each cancer type using the “rank_genes_groups” function in Scanpy. Genes were considered significant DEGs if they had an adjusted P-value < 0.05 and absolute log_2_(Fold Change) (|log_2_FC|)> 0.1. Because the BRCA data from phs002748 dataset contained only one tumor-reactive T cell, it was excluded from this differential expression analysis.

We used UpSet plots to visualize the intersection of DEGs across multiple cancer types. From this, we derived a final ccTR signature of 814 genes. This signature includes genes that showed consistent directionality (all Log_2_FC>0 or Log_2_FC<0) across all cancers and were significantly differentially expressed (adjusted p-value<0.05 and |Log_2_FC|>0.1) in at least one cancer. The ccTR signature is designed to be inclusive; rather than being restricted to significant genes in every cohort, it allows the machine learning model to learn the most predictive cross-cancer features during training.

For benchmarking, we included several previously reported marker genes (*CXCL13, ENTPD1*) and gene signatures (NeoTCR8, TR30, Caushi et al., Hanada et al.). To evaluate these signatures, we used the Scanpy “score_genes” function. This function calculates a single activity score per cell by taking the average expression of the signature genes and subtracting the average expression of a randomly sampled set of background genes.

To identify conserved biological processes, we performed overrepresentation analysis using the “enrichGO” function from the clusterProfiler (v4.12.3) package^39^. We focused on significant DEGs shared across multiple cancers to pinpoint common pathway shifts associated with tumor reactivity.

### Generative data augmentation

Significant imbalances in cell counts across cancer types and reactivity groups (reactive vs. non-reactive) initially led to poor model performance. Models trained on the original, imbalanced data were biased toward the majority class, predicting almost all query cells as non-tumor-reactive. To address this, we implemented a generative data augmentation strategy.

We utilized the scVI model (v1.4.0), which is based on a VAE architecture, for data augmentation^40^. We trained an individual scVI model for each dataset to learn the posterior predictive distribution of gene expression, denoted as 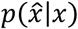. Here, *x* represents the input gene expression and 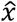 represents the sampled expression. Using the “posterior_predictive_sample” function from scVI, we generated synthetic cells that share the same transcriptomic features as the original populations.

Since tumor-reactive cells were the minority population, we specifically performed generative sampling for this group in the GSE176021, phs002748, and phs002765 datasets. Conversely, to prevent the NSCLC and HNSC cohorts (GSE176021 and GSE180268) from dominating the training process, we down-sampled these larger datasets. Following these adjustments, the proportion of tumor-reactive T cells was stabilized at approximately 30% across all training datasets. This threshold prevents model bias while remaining consistent with biological observations that reactive T cells represent a distinct minority of total TILs.

Importantly, augmented data were used exclusively for training the various machine learning architectures, including Turep, scANVI+scArches, and XGBoost. Performance evaluation was conducted solely on the original, non-augmented data. Compared to models trained on the raw datasets, those trained with generative augmentation showed a dramatic increase in predictive accuracy and robustness.

### Model design

Turep is built using the scvi-tools (v1.4.0) framework and is specifically optimized for cross-cancer prediction^40^. The framework consists of two components: Turep-sc, a pre-trained model for scRNA-seq data, and Turep-st, a model based on the FADVI architecture for ST data. We developed Turep-st because some ST platforms do not integrate well with scRNA-seq references in a shared latent space using Turep-sc. In these cases, using the standard Turep-sc model could lead to unreliable predictions.

Turep-sc is adapted from scANVI with a focal loss classification head for tumor reactivity labels^21,24^. An encoder maps the gene expression (*x*) to first latent variable (*z*_1_):

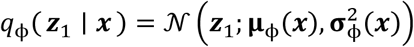

First latent variable is used for classifying tumor reactivity labels (*y*):

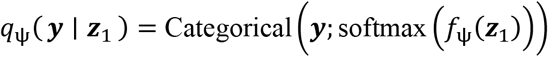

And for mapping to second latent variables (*z*_2_):

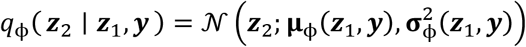

During generative process, first latent variable is mapped from *y* and *z*_2_:

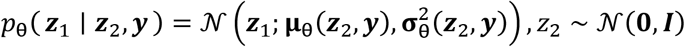

Gene expression is modeled as the zero-inflated negative binomial distribution (ZINB):

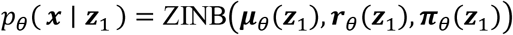

To better address the data imbalance issue, we used the supervised focal loss instead of regular cross-entropy loss for the classification:

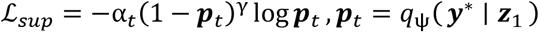

Where ***p***_*t*_ is the predicted probability of the true class *y*^*^, *α*_*t*_ is the class-balancing weight, and *γ* is the focusing parameter.

Total loss combines the supervised loss with unsupervised loss, which includes the KL divergence and reconstruction loss.

For scRNA-seq query datasets, Turep-sc employs the scArches strategy for transfer learning^22^. During this process, the model freezes specific components, including the first layers of the encoder and decoder, the encoder batch-normalization weights, and the classifier. The query and reference datasets are then trained together for 50 epochs to optimize data integration and ensure accurate label projection.

For prediction in ST data, Turep-st is adapted from FADVI to enhance data integration between scRNA-seq and ST data^34^. We separate the latent space in VAE into three subspaces capturing label-related (l), batch-related (b), and residual-related (r) variation.

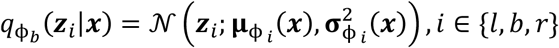

The latent variables from three subspaces are concatenated as decoder input:

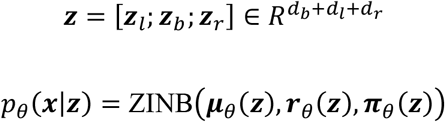

Classifiers use corresponding latent variables to predict cell labels and batches:

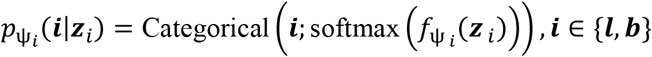

For cell label classification, we use the same focal loss as Turep-sc:

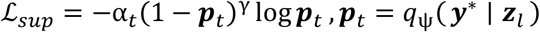

Adversarial classifier with gradient reversal (GR) layer is included to prevent label prediction from batch latent variable:

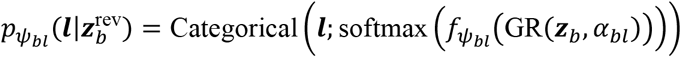

Other three classifiers are included to prevent batch prediction from label latent variable, and label or batch prediction from residual latent variable.

Total loss combines the KL divergence and reconstruction loss, supervised loss, adversarial loss, and cross-covariance loss.

### Performance evaluation

We used a “leave-one-cancer-out” cross-validation strategy to evaluate how well the model generalizes across different cancer types. In this approach, we trained the model on six cancer types and tested it on the seventh type. This process was repeated until every cancer had been used as a test set. We then calculated the mean metrics across all seven test iterations to measure how the model performs on cancers it has not seen before.

We treated the prediction of tumor-reactive T cells as a binary classification task, labeling tumor-reactive cells as positive and non-reactive cells as negative. Using scikit-learn (v1.5.1) and imbalanced-learn (v0.13.0), we measured performance with several metrics: accuracy, sensitivity, specificity, G-mean, AUROC, and average precision. Because our datasets were highly imbalanced, we focused on the G-mean and AUROC to identify the most robust and accurate methods.

### Benchmarking with other models

We included the following method for benchmarking cross-cancer tumor-reactive T cell prediction.

#### MaxGmean

This baseline uses a grid-search strategy to identify the classification threshold that maximizes the average G-mean. To implement this, we first calculated an activity score for each gene signature. We then generated a linearly spaced vector of 1,000 potential thresholds. Cells with an activity score above a given threshold were classified as tumor-reactive, while those below were classified as non-tumor-reactive. We calculated G-mean values for each of the six training cancers and selected the threshold that produced the highest average G-mean across those datasets.

#### XGBoost

We utilized XGBoost (v2.1.2), a robust gradient-boosting framework. This architecture is the basis for the existing predicTCR model used for reactivity prediction^25^. For each gene signature, we trained a model using the filtered gene expression matrix and the corresponding reactivity labels from six cancers. We found that models using log-normalized data outperformed those using raw counts; therefore, we used log-normalized input for our final XGBoost comparisons.

#### scANVI+scArches

We further benchmarked against scANVI and scArches, which currently represent the state-of-the-art methods for single-cell label projection. To train the scANVI model, we used raw counts as input, reactivity status as the cell label, and cancer type as the batch variable. Following standard protocols, we first trained the model for 50 epochs using scVI model, followed by 30 epochs using scANVI model. We then used scArches for transfer learning to expand the model to the query cancer types. During this stage, we trained for an additional 50 epochs while freezing the first layers of the encoder and decoder, the encoder batch-normalization weights, and the classifier.

For the XGBoost and scANVI+scArches frameworks, we independently trained multiple models using different gene signatures as input features. We then compared the testing performance of these configurations to select the best-performing model for each architecture. Notably, although we requested access to the original predicTCR model, the weights were not shared by the authors. Consequently, we were unable to include the specific pre-trained predicTCR model in our benchmarking and instead relied on our independent XGBoost implementation.

### Feature attribution analysis

To identify the biological drivers behind Turep’s predictions, we performed feature attribution analysis using the final Turep-sc model trained on all cancer datasets. This analysis allows us to move beyond black-box classifications and understand which specific genes most strongly influence the model’s decisions.

We implemented feature attribution in both the Turep-sc and Turep-st models using the Captum library (v0.7.0). Specifically, we used the GradientShap class to measure gene importance. GradientShap is a gradient-based method that approximates SHAP values. These values are rooted in cooperative game theory and ensure that the credit for a prediction is distributed fairly among the input genes. As a robust alternative, we also implemented the IntegratedGradients method within our framework.

When the model makes a prediction, it calculates a SHAP value for every gene in each individual cell. This provides a local explanation, showing how each gene contributes to that specific cell’s classification. A positive SHAP value indicates that the gene’s expression pushes the model toward a “tumor-reactive” prediction. Conversely, a negative SHAP value suggests that the gene’s expression pushes the model toward a “non-tumor-reactive” label.

To aggregate these cell-specific explanations to a global understanding of the model, we calculated the mean and standard error of the SHAP values across the entire dataset. By ranking genes according to their mean attribution scores, we identified the most critical positive and negative contributors to tumor reactivity. This approach allows us to determine if certain genes serve as universal markers of reactivity across all cancers or if their importance is specific to certain malignancies.

### Turep application in independent scRNA-seq datasets

We used several independent scRNA-seq datasets to investigate the relationship between tumor-reactive T cells and therapy response. These included: BCC patients with anti-PD1 therapy (GSE123814)^29^, BRCA patients with anti-PD1 therapy and chemotherapy (BioKey)^27^, BRCA patients with anti-PD1 therapy and radiotherapy (GSE246613)^28^, RCC patients with anti-PD1 therapy (SCP1288)^30^, SKCM patients with ACT therapy (GSE229861)^31^.

We applied the pre-trained Turep model to these query datasets using the scArches transfer learning strategy. To minimize batch effects between samples, Turep generated predictions for each sample individually before the results were concatenated. For each sample, we calculated the percentage of CD8^+^ T cells predicted to be tumor-reactive. Based on our results and previous studies of TILs, we used a threshold of 10% or higher to indicate a strong anti-tumor potential.

We grouped samples based on their treatment types (ICB, ACT, chemotherapy, or radiotherapy) and their clinical responses as defined in the original studies. We categorized “Good responses” as samples labeled as Response (R), Partial Response (PR), Complete Response (CR), or significant T cell clonal expansion (E). We categorized “Poor responses” as Non-responder (NR), Stable Disease (SD), Progressive Disease (PD), or limited T cell clonal expansion (NE). We compared the percentages of tumor-reactive T cells between these groups using the Wilcoxon test. For longitudinal datasets, we treated samples from the same patient as paired samples.

Finally, we leveraged dataset GSE283942 to study the physical interactions between tumor-reactive T cells and their targets^32^. This dataset is unique because it contains both single cells (singlets) and small cell clusters captured together (doublets or multiplets). Because these groups were sorted into different samples for sequencing, we could distinguish them in the merged data. We used Turep to predict the reactivity of both singlets and multi-cell clusters that contained T cells. However, we noted a potential limitation: because Turep was trained specifically on singlet T cells, its predictions for doublets or multiplets might be less accurate if the gene expression profile is dominated by the interacting tumor cell or APC.

### Turep application in independent ST dataset

We used the SPATCH dataset to investigate the spatial distribution of tumor-reactive T cells^33^. This dataset includes three cancer tissues profiled across four ST technologies: Visium HD (Fresh Frozen, FF or Formalin-Fixed Paraffin-Embedded, FFPE), Stereo-seq, Xenium, CosMx. We selected CD8^+^ T cells for analysis based on the cell-level or 8-μm spot-level (Visium HD) original annotations provided by the authors.

To perform the predictions, we trained a Turep-st model for each individual ST sample. We observed that the expression profiles of single-cell and spatial data differed significantly, which resulted in predicted tumor-reactive probabilities consistently below 0.6. To account for this modality gap and ensure more reliable identification, we calculate tumor-reactivity percentile (*P*_*i*_) from the rank of probability (*R*_*i*_):

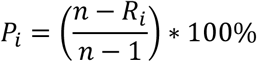

Using this metric, we categorized CD8^+^ T cells into three groups: Top 10% (*P*_*i*_ ≥ 90%, high-confidence reactive), 10%∼50% (50% ≤ *P*_*i*_ < 90%, potentially reactive), Low 50% (*P*_*i*_ < 50%, high-confidence non-reactive). This stratification allowed us to compare the biological niches of T cells with varying reactive potential regardless of the absolute probability scores.

We characterized the spatial microenvironment of these T cells using Squidpy (v1.6.5)^41^. For each CD8^+^ T cell, we identified its 20 nearest spatial neighbors and analyzed their cell-type composition. To validate the functional activity of the predicted cells, we examined the expression of key genes involved in cell-mediated immune responses, including *B2M, GZMB, GZMK*, and *PRF1*, within these neighbor populations. We calculated the average expression of these genes across neighbor cell types and used the Wilcoxon rank-sum test to compare the mean neighbor expression levels between the different T cell groups.

Finally, we identified active tumor-recognition regions by locating areas where high T cell tumor reactivity percentiles overlapped with elevated neighbor B2M expression. Using the SpatialData framework (v0.4.0)^42^, we extracted 1000×1000 pixel regions centered on high-probability tumor-reactive T cells. These regions typically contained clusters of CD8^+^ T cells, APCs, and tumor cells, resembling previously reported CRATER niches^36^. This spatial convergence provides independent evidence that Turep-st correctly identifies T cells that are actively engaged in antigen recognition within the tissue architecture.

## Data availability

This study used published datasets for investigation and machine learning model training. The paired scRNA-seq and scTCR-seq datasets for model training include publicly available GEO datasets GSE176021, GSE180268, and controlled accessed dbGaP datasets phs001451, phs002748, phs002765. The scRNA-seq datasets demonstrating Turep application are from publicly available, including GSE123814, GSE229861, GSE246613, GSE283942 from GEO database; SCP1288 from Single Cell Portal; and https://lambrechtslab.sites.vib.be/en/single-cell (BioKey). The ST dataset is available at https://spatch.pku-genomics.org/#/download.

## Code availability

Turep code is publicly available at https://github.com/liuwd15/turep, and the pretrained model is available at https://zenodo.org/records/19005962.

## Acknowledgements

This work was partially supported by National Institutes of Health grants R01LM012806 and U01AG079847. We thank the technical support of the Cancer Genomics Core supported by the Cancer Prevention & Research Institute of Texas (CPRIT) grant RP240610. W.L. was supported by John and Rebekah Harper Fellowship in Biomedical Sciences and CPRIT Fellowship in the Biomedical Informatics, Genomics and Translational Cancer Research Training Program (BIG-TCR, CPRIT RP210045). The funders had no role in the study design, data collection and analysis, decision to publish or prepare the manuscript.

## Author contributions

W.L: Conceptualization, Methodology, Software, Formal analysis, Investigation, Data Curation, Writing - Original Draft, Writing - Review & Editing. C.T.: Investigation, Data Curation, Writing - Review & Editing. E.M.S.: Writing - Original Draft, Writing - Review & Editing. Z.Z.: Writing - Original Draft, Writing - Review & Editing, Supervision, Funding acquisition.

## Declaration of interests

The authors declare no competing interests.

## Notes

### Competing Interest Statement

The authors have declared no competing interest.

