## Supplementary Fig for "Turep: Detecting cross-cancer tumor-reactive T cells in single-cell and spatial transcriptomics data"

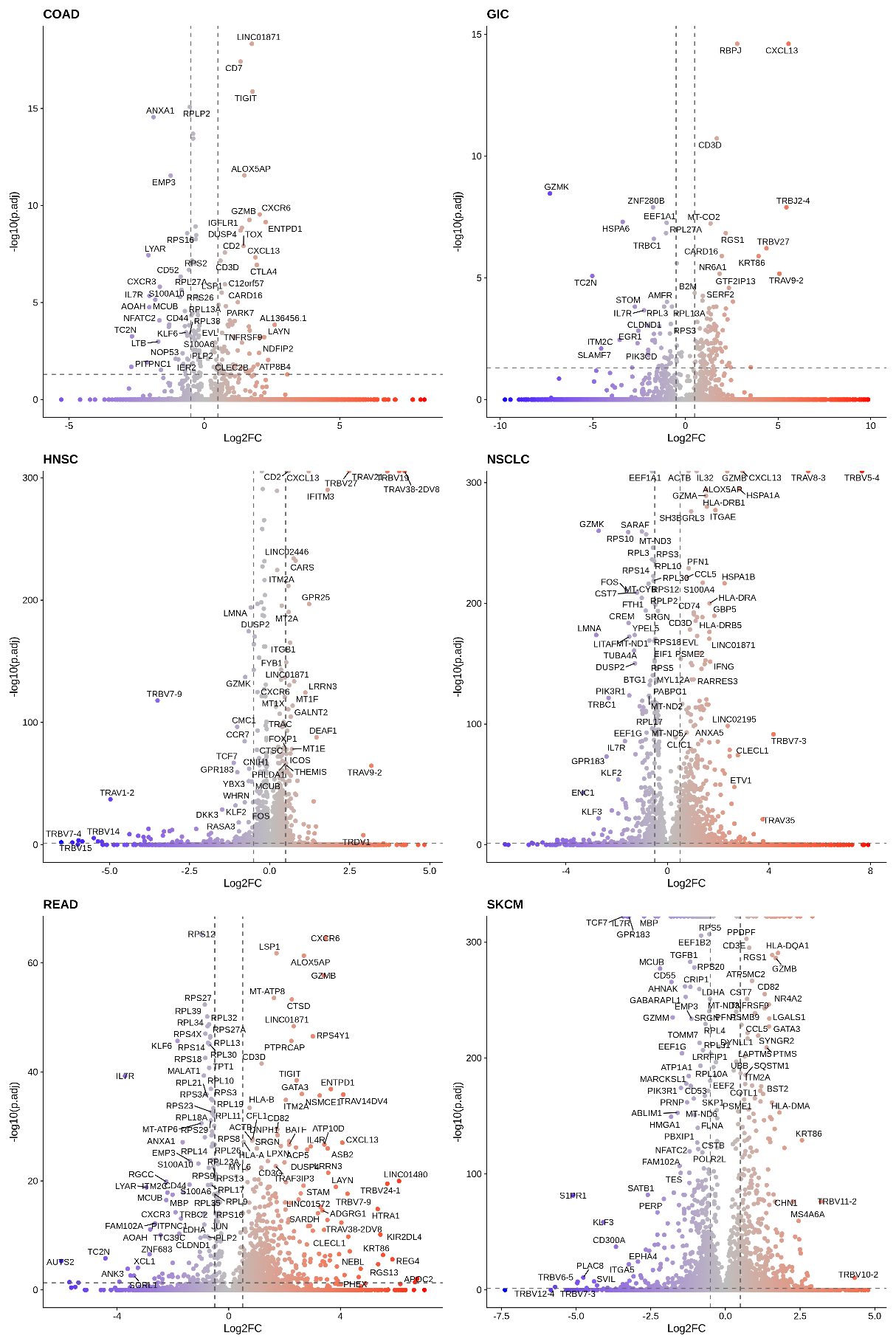

**Supplementary Fig. S1. Volcano plots showing differential expressed genes (DEGs) between tumor-reactive T cells and non-tumor-reactive T cells from each cancer.**

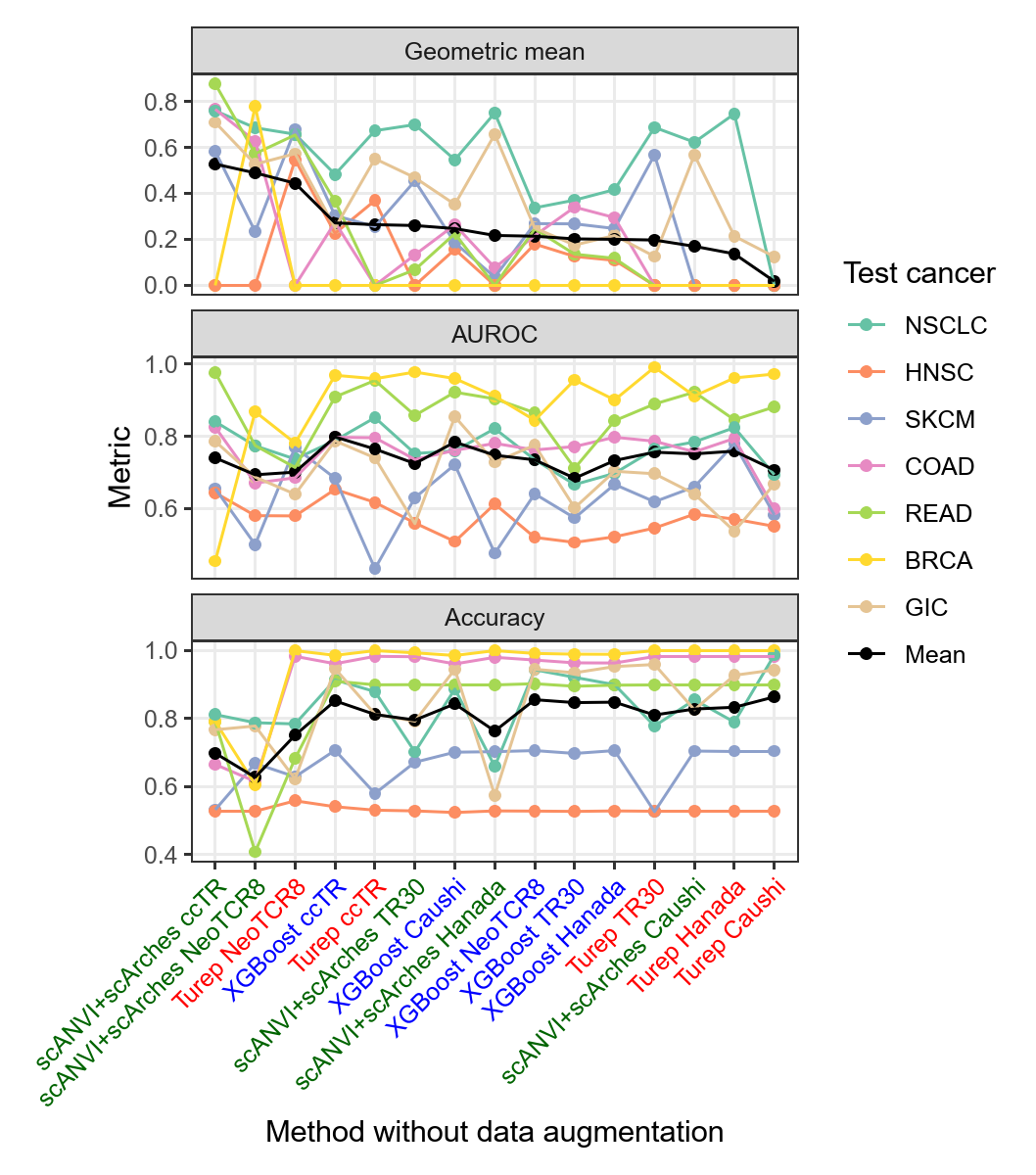

**Supplementary Fig. S2. Performance of machine learning models without generative data augmentation.**

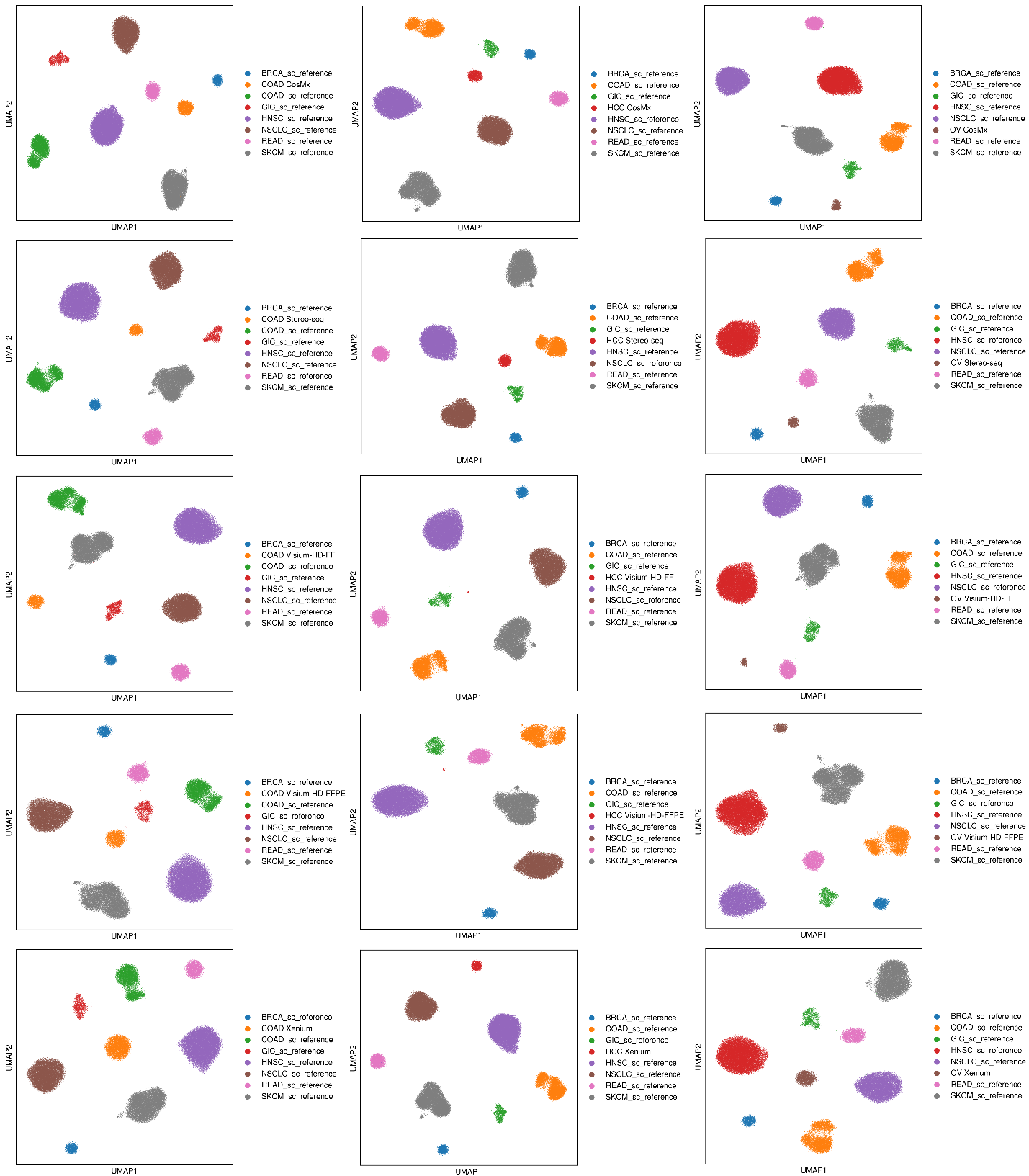

**Supplementary Fig. S3. UMAP plots of cell representations in batch subspace from Turep-st.** When making prediction in each ST sample, the batch subspace captures variations of technologies and cancer types, showing as discrete clusters.

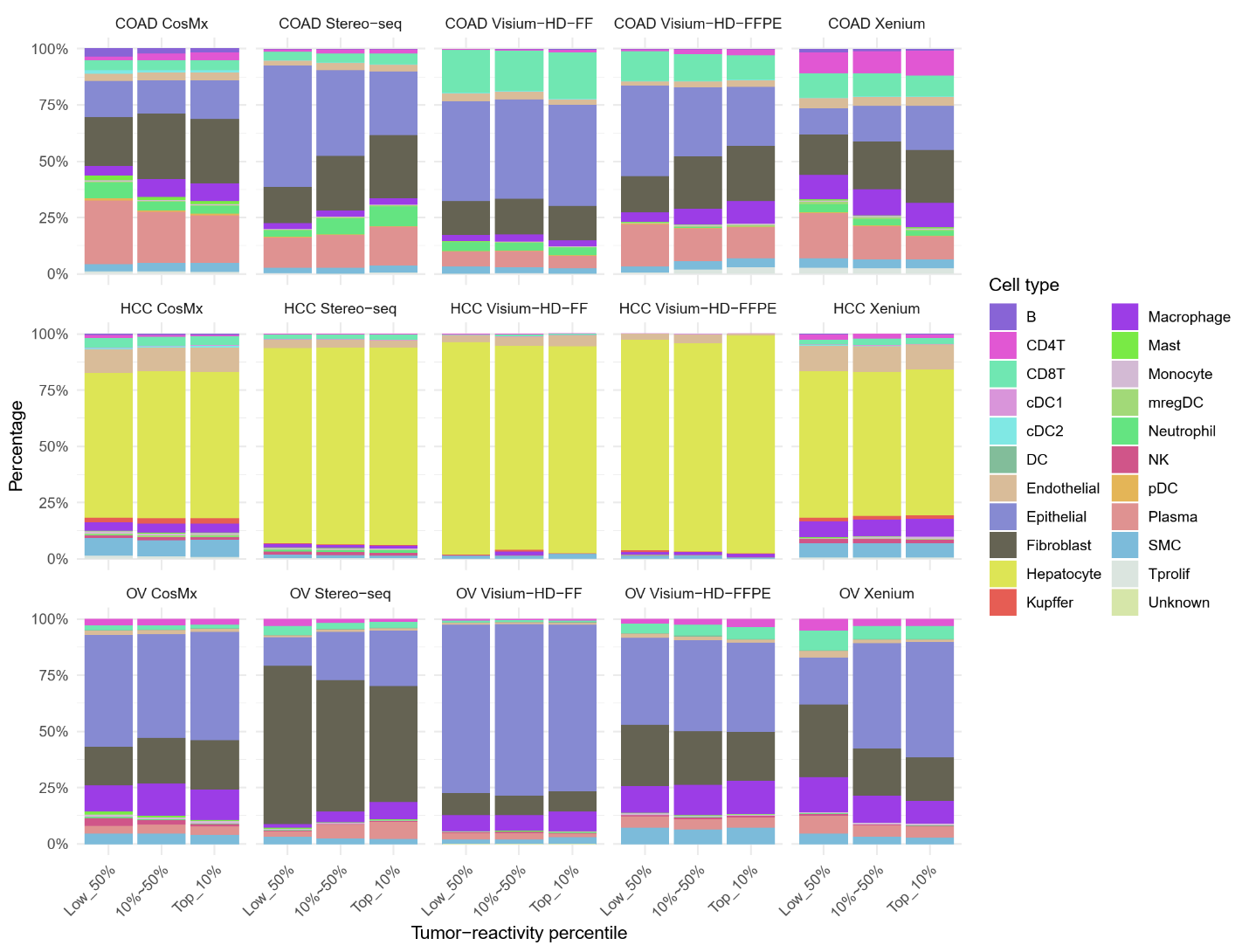

**Supplementary Fig. S4. Neighbor cell compositions of CD8^+^ T cells in different reactivity percentile groups.** There is a strong variation in neighbor cell compositions of CD8^+^ T cells across different ST technologies.

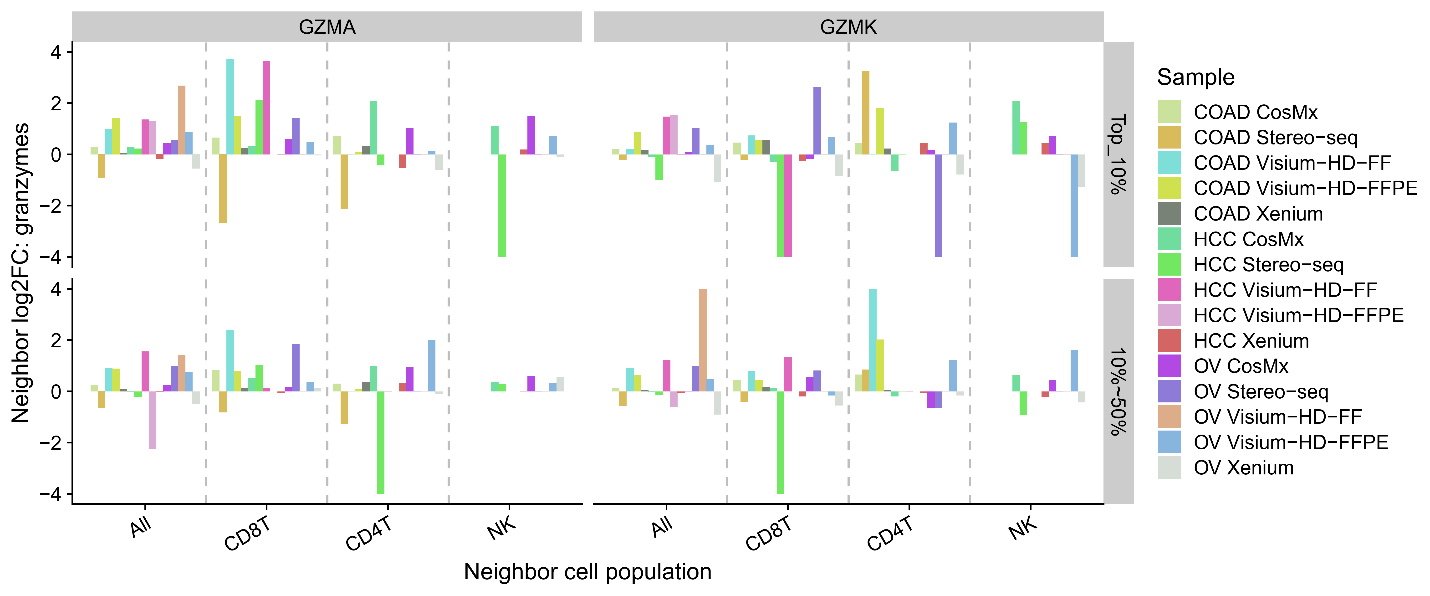

**Supplementary Fig. S5. Differential expression analysis of GZMA and GZMK in neighboring cell populations.** Bar plots represent the log2(Fold Change) in neighbors of likely tumor-reactive T cells (top 10% or 10%~50% probability) compared to bottom 50% probability T cells across multiple ST datasets.
